# Transection injury differentially alters the proteome of the human sural nerve

**DOI:** 10.1101/2021.11.23.469670

**Authors:** Monica J. Chau, Jorge E. Quintero, Eric Blalock, Christopher Samaan, Greg Gerhardt, Craig van Horne

**Affiliations:** Brain Restoration Center, College of Medicine, University of Kentucky, Lexington, KY, United States of America; Department of Neurosurgery, College of Medicine, University of Kentucky, Lexington, KY, United States of America; Department of Neuroscience, College of Medicine, University of Kentucky, Lexington, KY, United States of America; Department of Pharmacology and Nutritional Sciences, College of Medicine, University of Kentucky, Lexington, KY, United States of America; Department of Neurology, College of Medicine, University of Kentucky, Lexington, KY, United States of America

**Author notes:** Corresponding author; (CV).

**Keywords:** Peripheral nerve repair, regeneration, peripheral nerve injury

## Abstract

Regeneration after severe peripheral nerve injury is often poor. Knowledge of human nerve regeneration and the growth microenvironment is greatly lacking. We aimed to identify the regenerative proteins in human peripheral nerve by comparing the proteome before and after a transection injury. In a unique study design, we collected from the same participants, samples from naïve and degenerating sural nerve. Naïve and degenerating (two weeks after injury) samples were analyzed using mass spectrometry and immunoassays. Using a correlation matrix, we found significantly altered levels following the nerve injury. Mass spectrometry revealed that post-injury samples had 672 proteins significantly upregulated and 661 significantly downregulated compared to naïve samples (q < 0.05, |FC| > 2). We used Gene Ontology pathways to highlight groups of proteins that were significantly upregulated or downregulated with injury-induced degeneration and regeneration. Significant protein changes in key pathways were identified including growth factor levels, Schwann cell de-differentiation, myelination downregulation, epithelial-mesenchymal transition, and axonal regeneration pathways. Having proteome signatures of human peripheral nerves of both the uninjured and the degenerating/regenerating state may serve as biomarkers to aid in the future development of repair strategies and in monitoring neural tissue regeneration.

## Introduction

After severe peripheral nerve injury, there is poor regeneration and functional recovery. Understanding the transcriptome[1], proteome, and the mechanisms that underlie peripheral nerve regeneration is essential to further develop peripheral nervous system (PNS) regenerative therapies and to improve clinical outcomes. Even though there is a great deal that is known about peripheral nerve regeneration and the growth-permissive microenvironment in animal models, results on human nerves is sparse and inconclusive[2,3]. Multiple components contribute to peripheral nerve repair after injury notably Schwann cells[4–7], fibroblasts, endothelial cells, and immune cells such as macrophages[8–10]. An injured peripheral nerve undergoes three main processes for recovery and reestablishment of a functional connection with its distal end[11]. First, within 24-48 hours after transection injury of the nerve, Wallerian degeneration starts in which the distal end of the transection undergoes axonal degeneration. Afterward, the myelin is degraded as macrophages infiltrate this area. Secondly, axonal regeneration begins, leading to the third step of end-organ re-innervation. Macrophages and Schwann cells play major roles in mediating degeneration and regeneration.

Animal models of peripheral nerve regeneration demonstrate that Schwann cells have remarkable plasticity. They rapidly adapt to injury through extensive cellular reprogramming that transforms their myelinating phenotype into a reparative phenotype[12,13]. Studies have delineated two phases of the Schwann cell injury response: 1) clearing of myelin[4] and de-differentiation of the cell’s myelinating phenotype and 2) maturation into the repair Schwann cell phenotype in which the cells promote survival by releasing neuroprotective factors and increasing anti-apoptosis factors. Schwann cells upregulate and release a whole host of neurotrophic and cell survival factors including glial cell-derived neurotrophic factor [14,15], neurotrophin-3 (NT-3), nerve growth factor (NGF)[16,17], brain derived neurotrophic factor (BDNF)[18], vascular endothelial growth factor (VEGF), erythropoietin [19], pleiotrophin, and N-cadherin[6,7,20–22].

Human peripheral nerve injury studies are typically limited to collecting nerve tissues *after* an injury (*e.g*. after a trauma) while obtaining healthy, uninjured nerve beforehand has not been as feasible. As such, many aspects of human peripheral nerve regeneration are unknown including the profile of protein upregulation and downregulation. Understanding these changes would help optimize discovery and clinical management strategies for improving PNS repair outcomes.

To address these gaps in knowledge, we performed a comprehensive proteomic analysis to include more than 5000 of the most abundant proteins found in peripheral (sural) nerve *before and after* a transection injury. We hypothesized that transecting a peripheral nerve and leaving it in place for two weeks would increase synthesis in the pathways associated with nerve regeneration. Thus, this proteome profile would provide a database to identify new targets for improving regeneration outcomes.

## Materials and Methods

### Research Subjects

The University of Kentucky’s Institutional Review Board approved the study and the participants provided written informed consent. Sural nerve tissue was collected from 26 participants before and after sural nerve transection *in situ*. Participants were diagnosed with idiopathic PD. Nine were female, 17 were male. Their ages ranged from 50-70 years old (mean 61 years old). Years diagnosed with PD ranged from 2-17 years (mean 10 years). Tables 1 and 2 list participant demographics from both types of proteomic analyses (mass spectrometry and targeted immunoassays). Note that there are three participants whose tissues were used in both mass spectrometry and targeted immunoassay.

**Table 1.** Mass Spectrometry Participant Demographics. Mass spectrometry study participant demographics listed include age, years diagnosed with PD, and assigned birth sex.

| **Participant** | **Age (years)** | **Years diagnosed** | **Assigned Birth Sex** |
| --- | --- | --- | --- |
| 1 | 70 | 7 | Male |
| 2 | 70 | 12 | Female |
| 3 | 63 | 11 | Male |
| 4 | 69 | 9 | Male |
| 5 | 66 | 2 | Female |
| 6 | 66 | 16 | Male |
| 7 | 52 | 6 | Female |
| 8 | 60 | 6 | Male |
| 9 | 57 | 10 | Male |
| 10 | 60 | 7 | Female |
| 11 | 50 | 8 | Male |
| 12 | 61 | 9 | Male |
| 13 | 61 | 8 | Male |
| 14 | 65 | 6 | Male |

**Table 2.** Targeted Immunoassays Participant Demographics. Immunoassay study participant demographics listed include age, years diagnosed with PD, and assigned birth sex.

| **Participant** | **Age (years)** | **Years diagnosed** | **Assigned Birth Sex** |
| --- | --- | --- | --- |
| 15 | 51 | 17 | Male |
| 7 | 52 | 6 | Female |
| 17 | 54 | 16 | Male |
| 18 | 56 | 10 | Male |
| 19 | 58 | 12 | Female |
| 20 | 60 | 13 | Female |
| 21 | 61 | 15 | Female |
| 13 | 61 | 8 | Male |
| 23 | 62 | 4 | Male |
| 24 | 62 | 7 | Male |
| 25 | 64 | 16 | Male |
| 26 | 65 | 8 | Male |
| 14 | 65 | 6 | Male |
| 28 | 66 | 9 | Female |
| 29 | 69 | 8 | Female |

### Peripheral Nerve Transection and Tissue Collection

Our approach in transecting the peripheral nerve and collecting tissue has been previously described[1,23,24]. Fig 1 illustrates the timing of the transection and tissue collections. Briefly, sural nerve samples were collected at two different time points: a naïve non-injured sample collected during the first stage of surgery followed by a post-transection (injured) sample from the second stage of surgery two weeks later (. To collect the naive tissue, the neurosurgeon identified the neurovascular bundle containing the sural nerve in the ankle. Two sutures were tied around the nerve, 1 cm apart. The section of nerve just proximal to the sutures was transected, and a 1-cm segment was excised for analysis. Individual nerve fascicles were separated, snap-frozen, and stored for assays as in Welleford et al.[1].

**Fig 1.**
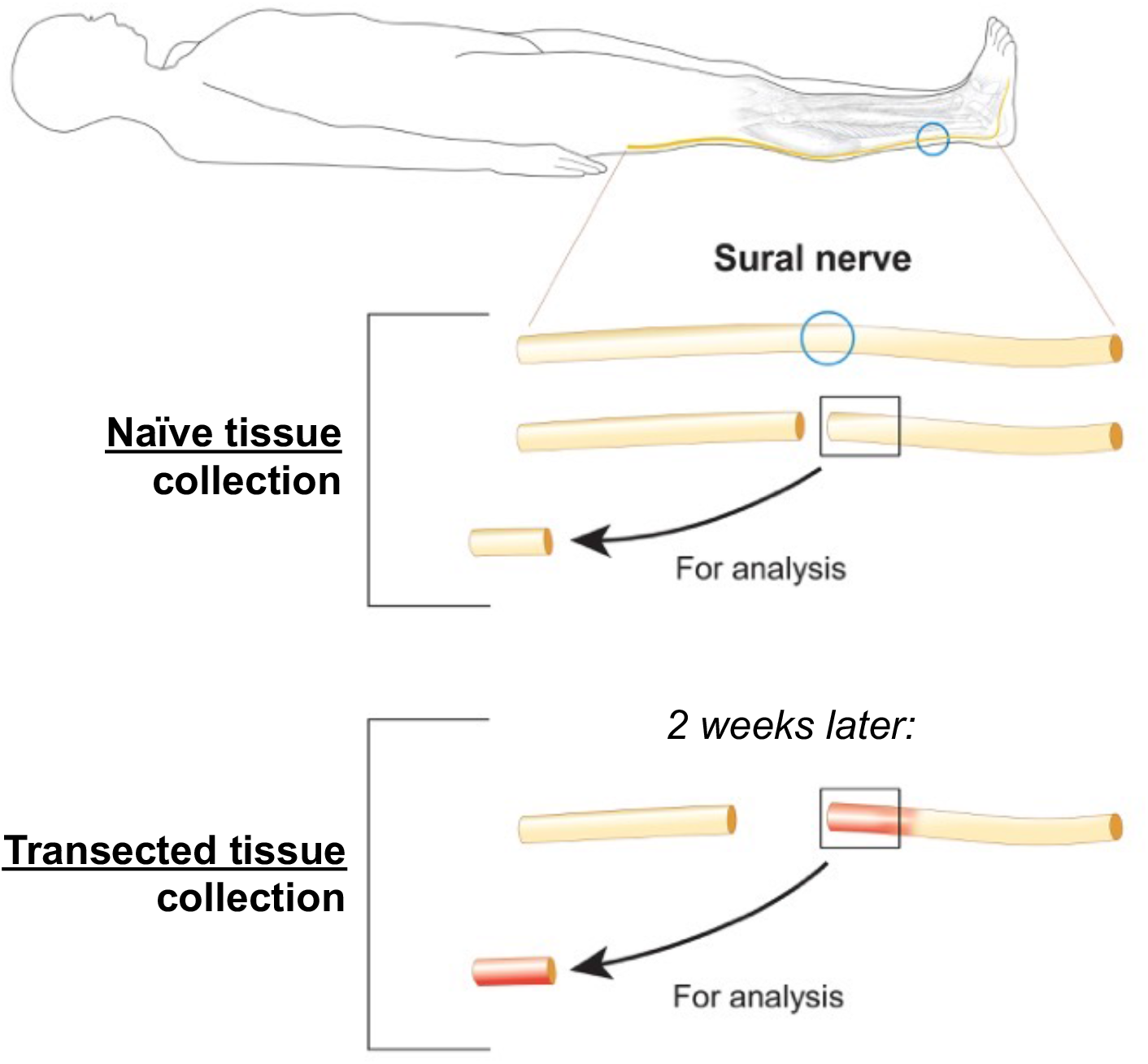
Study Overview. Schematic to illustrate the timing and location of the sural nerve injury and tissue collection

To collect post-transection tissue for both implantation in the clinical trial and proteomic analysis, the ankle incision was reopened after two weeks, suture markers were located, and a new 1-2 cm segment was collected from the distal nerve stump for analyses and implantation as part of the trial design. Individual nerve fascicles were separated, snap-frozen, and stored for assays[1].

### Proteomics Analyses

#### Mass spectrometry proteomics

TMT analysis was used which is a global, unbiased method. As many proteins/peptides as possible were detected and searched against a species-specific database. Total protein was isolated from peripheral nerve tissue samples. Protein samples were prepared for mass spectrometric analysis, and MS/MS sequence and tandem mass tag reporter ion data was collected using a state-of-the-art Orbitrap Eclipse instrument. Quantitative analysis was performed to obtain a comprehensive proteomic profile.

Proteins were identified and quantified using MaxQuant and visualized with Scaffold using 1% false discovery thresholds at both the protein and peptide level. The data was checked for quality and normalized using an in-house ProteiNorm Shiny app, a tool for a systematic evaluation of normalization methods, imputation of missing values and comparisons of different differential abundance methods.

##### CME bHPLC TMT Methods – Orbitrap Eclipse

Proteins were reduced, alkylated, and purified by chloroform/methanol extraction prior to digestion with sequencing grade modified porcine trypsin (Promega). Tryptic peptides were labeled using tandem mass tag isobaric labeling reagents (Thermo) following the manufacturer’s instructions and combined into three 11-plex sample groups with a pooled reference sample in each group. Each labeled peptide multiplex was separated into 46 fractions on a 100 x 1.0 mm Acquity BEH C18 column using an UltiMate 3000 UHPLC system (Thermo) with a 50 min gradient from 99:1 to 60:40 buffer A:B ratio under basic pH conditions, and then consolidated into 18 super-fractions. Each super-fraction was then further separated by reverse phase XSelect CSH C18 2.5 μm resin on an in-line 150 x 0.075 mm column using an UltiMate 3000 RSLCnano system (Thermo). Peptides were eluted using a 60 min gradient from 98:2 to 60:40 buffer A:B ratio. Eluted peptides were ionized by electrospray (2.2 kV) followed by mass spectrometric analysis on an Orbitrap Eclipse Tribrid mass spectrometer (Thermo) using multi-notch MS3 parameters with real-time search enabled. MS data were acquired using the FTMS analyzer in top-speed profile mode at a resolution of 120,000 over a range of 375 to 1500 m/z. Following CID activation with normalized collision energy of 35.0, MS/MS data were acquired using the ion trap analyzer in centroid mode and normal mass range. Using synchronous precursor selection, up to 10 MS/MS precursors were selected for HCD activation with normalized collision energy of 65.0, followed by acquisition of MS3 reporter ion data using the FTMS analyzer in profile mode at a resolution of 50,000 over a range of 100-500 m/z.

Buffer A = 0.1% formic acid, 0.5% acetonitrile
Buffer B = 0.1% formic acid, 99.9% acetonitrile
Both buffers adjusted to pH 10 with ammonium hydroxide for offline separation

##### Mass Spectroscopy Statistical Analysis

Each protein in the data set had a log fold change and false discovery rate (FDR) adjusted p-value. For the proposed experiments, the units used for statistical comparison were normalized MS3 TMT reporter ion intensities. We used the limma package for statistical analysis, which is a moderated t-test that accounts for the data variance. We ran a paired sample model using limma. The final results contained the log2 fold change, unadjusted p-values, and FDR adjusted p-values. Significance threshold is a Fold change > 2 and a FDR adjusted p-value < 0.05. Each data set had its own characteristics so we performed QC checks of data prior to running statistical analyses to ensure the most appropriate normalization method and statistical test was run.

##### Correlation matrix

Mass spectrometry quantification (log 2 scale, Cyclic Loess normalized) data included 5,573 rows for different proteins mapped back to their gene symbol representations for 28 paired (naïve and injured) samples from 14 subjects.

##### Pre-statistical procedures

###### ‘0’ values

Seventeen percent of the data space (# rows * # samples = 156,044) was occupied by ‘0’ values. These ‘0’ values could represent low levels (below the detection threshold), no levels, or technical error[25]. In proteomic data, ‘0’s are considered missing-not-at-random’ (MNAR), and various strategies have been proposed to address this property of the data[26]. Here, we used a two-step procedure to address the ‘0’s. First, the number of ‘0’s on each row were quantified and rows with < 20 / 28 ‘0’s (i.e., at least seven reliable signal intensity measures on the row) were retained for further analysis (5,508 rows of data were retained). Second, within this filtered data set, the remaining 0’s were treated as missing values.

###### Annotation

Fourteen protein symbols with date-like names (from the MAR, MARC, SEP, and SEPT families) were updated to reflect newer non-date-like protein symbols (e.g., SEP15 was changed to SELENOF) to avoid their accidental conversion to date codes in spreadsheet programs. Additionally, 434 instances of ‘repeated’ annotations (that is, more than one row reporting the same gene symbol) were identified. Among these, the repeated protein symbol with the highest average level was retained for analysis, resulting in a final data set of (28 samples * 5074) uniquely protein symbol mapped proteins with reliable signal intensities.

##### Volcano plot

For the total of 5074 proteins included in the analysis, log 2 fold changes (x axis) were plotted against the log 10 p-values (y axis) for each protein. Highly stringent p-value (≤ 1E-10) and fold change (FC) (≥ |16|; or as expressed on the log 2 scale, ≥ |4|) cutoffs were used to highlight proteins with the largest and most significant changes.

##### GO pathway analyses and heatmap generation

Gene ontology pathways that were specifically related to peripheral nerve injury and regenerative processes were analyzed. The visualized mapped proteins represent the significantly differentially expressed proteins from each pathway. The proteins that corresponded to the GO pathways that were significantly upregulated or downregulated based off the p value and Log 2 FC were graphed in clustered hierarchical heatmaps using Ward’s hierarchical clustering.

Heatmaps were created in JMP software (SAS Institute, Cary NC). The protein and its fold change were mapped using hierarchical clustering. The range of fold changes were illustrated with a red/blue gradient and 0 was set at the center with gray. Note that participants’ samples that lacked either naïve or injured tissue results (e.g. because of insufficient amounts of tissue) were removed from analysis so as to not misrepresent fold changes. These are marked with a black and white grid pattern in the heatmaps.

#### Immunoassays for proteins of interest: ELISA and Luminex

Certain proteins of interest were not detected because they were not in high enough abundance to be detected by mass spectrometry. For these proteins of interest, such as growth factors, we used enzyme-linked immunosorbent assay (ELISA) and multiplex Luminex^®^ immunoassays (Cincinnati Children’s Hospital Flow Cytometry Core) to detect their change after injury. We compared naïve and injured tissues, from 15 participants, three of these participants’ tissues were also used in mass spectrometry analysis (see Tables 1 and 2).

Analyte concentrations in the sample supernatants were determined by ELISA according manufacturer’s protocol. The ELISA and antibody sources and dilutions were: Cerebral dopamine neurotrophic factor (CDNF) (Abcam, Cambridge, MA), samples were diluted 1:100. Nuclear factor erythroid 2-related factor 2 (NRF2) (ThermoFisher Scientific, Carlsbad, CA), samples diluted 1:2. Nerve Growth Factor Receptor (NGFR) (ThermoFisher Scientific, Carlsbad, CA), samples were diluted 1:2. B cell lymphoma 6 (BCL-6) (MyBiosource, San Diego, CA), samples were neat. EPO concentrations in the sample supernatants were determined by using MilliplexTM Multiplex kits (MilliporeSigma, Darmstadt, Germany) and BDNF, Beta-NGF, Platelet-Derived Growth Factor (PDGF)-AA, VEGF, GDNF, NT-3, PDGF-BB, PDGF-AB were determined by Human Magnetic Luminex Assays (R&D Systems, Minneapolis, MN) according to manufacturer’s protocol.

##### Analysis

For each protein analyzed with the immunoassay, the mean difference between the naïve and injured samples and the 95% confidence interval (Excel, Microsoft) is reported.

#### Availability of Data and Materials

Datasets are available at (https://uknowledge.uky.edu/).

## Results

### A clear proteomic distinction between naïve and injured nerve

Protein levels were significantly altered after nerve injury (Fig 2A) based on Pearson’s *r* correlation. The Pearson’s correlation r-value for each sample, correlated with every other sample is displayed in a correlation matrix.

**Fig 2.**
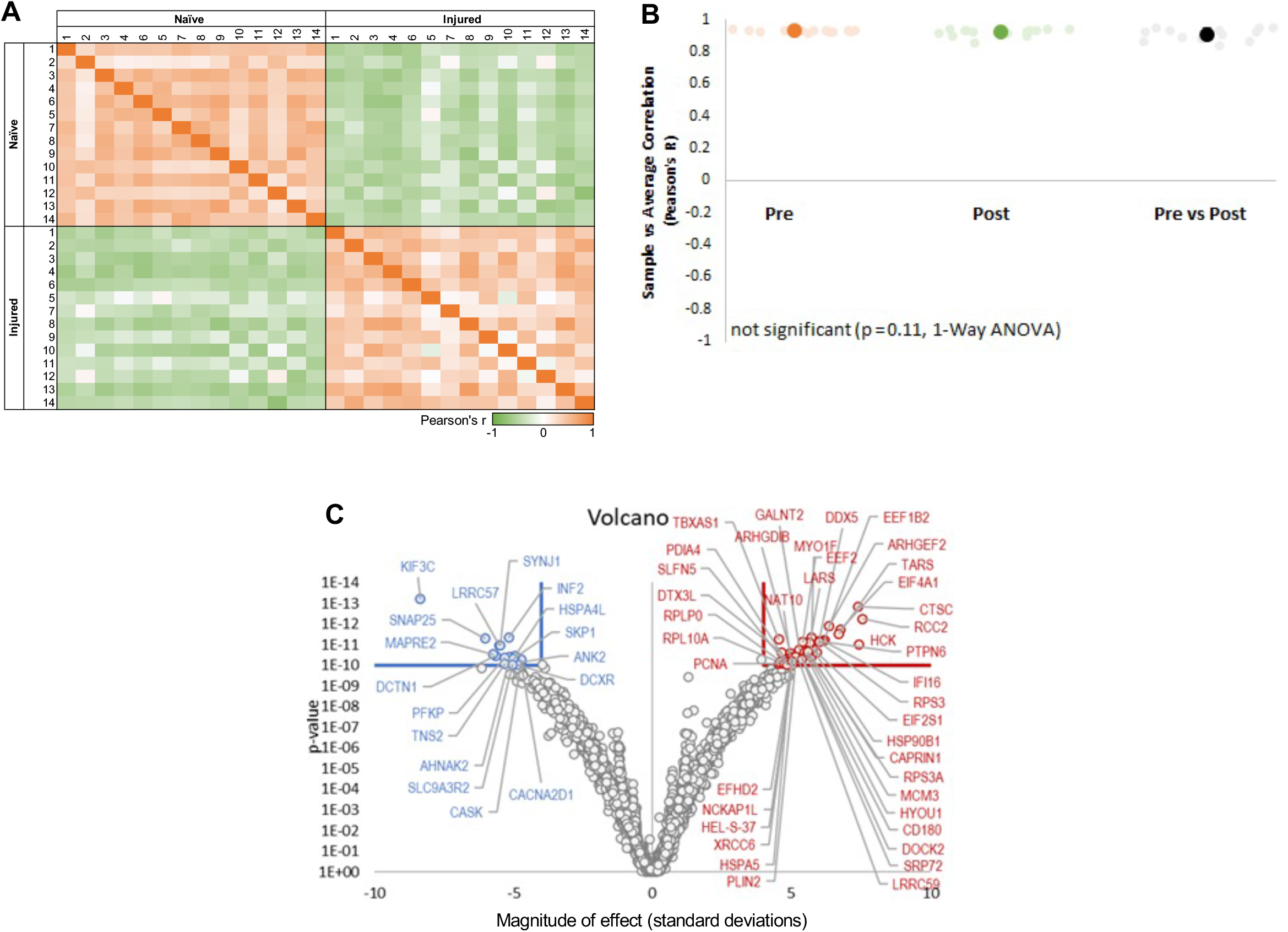
Clear Proteomic Changes Between Naïve and Injured Nerve. **A.** Protein levels were significantly altered after nerve injury based on Pearson *r* correlation. The Pearson’s correlation R-value for each sample, correlated with every other sample is displayed in a correlation matrix of naïve and injured samples (matrix scale orange R = 1: more similar, green R = −1: less similar). **B.** Lin’s CCC demonstrated strong homogeneity among samples within the pre- and post-injury groups **C.** The volcano plot shows proteins that were downregulated (blue) and upregulated (red) after injury

We calculated Lin’s Concordance Correlation Coefficient (CCC), which quantifies degree of similarity among different samples. Results indicated strong positive, and highly similar profiles from each sample compared to the average of all samples within either the pre-injury or post-injury groups (Fig 2B). CCCs in Pre-injury (CCC= 0.936 +/- 0.0065) and Post-injury (CCC = 0.93 +/- 0.012) conditions were greater than the 0.8 typically interpreted as showing strong similarity, and thus indicates generally strong homogeneity among samples within the pre- and post-injury groups.

Among the largest and most significant protein synthesis changes of the *whole* mass spectrometry dataset, the volcano plot shows 17 proteins that were downregulated (blue) and 41 upregulated proteins (red) after injury (Fig 2C). The volcano plot cutoffs were more stringent than in each individual Gene Ontology (GO) heatmap and highlight the most upregulated and downregulated proteins of the whole data set.

### Significant changes in protein changes shown in Gene Ontology (GO) pathways and heatmaps

Mass spectrometry analysis of naïve and injured sural nerve returned a total of 5573 of the most abundant proteins in the tissues. When comparing naïve to injured nerve, 672 proteins were significantly upregulated and 661 were significantly downregulated. We used heatmaps to highlight significantly changed GO pathways related to tissue injury degeneration and regeneration.

#### Growth Factors

##### Mass Spectrometry

Fig 3 shows all of the proteins that were significantly upregulated or downregulated (*q* < 0.05, |FC| > 2) from the GO pathway, Growth Factor Activity (GO:0008083). Six proteins were significantly downregulated after transection, and five proteins were significantly upregulated after injury (Fig 3). Out of 163 unique genes listed in this GO pathway, 11 corresponding proteins (7%) were significantly differentially expressed when comparing naïve and injured tissue.

**Fig 3.**
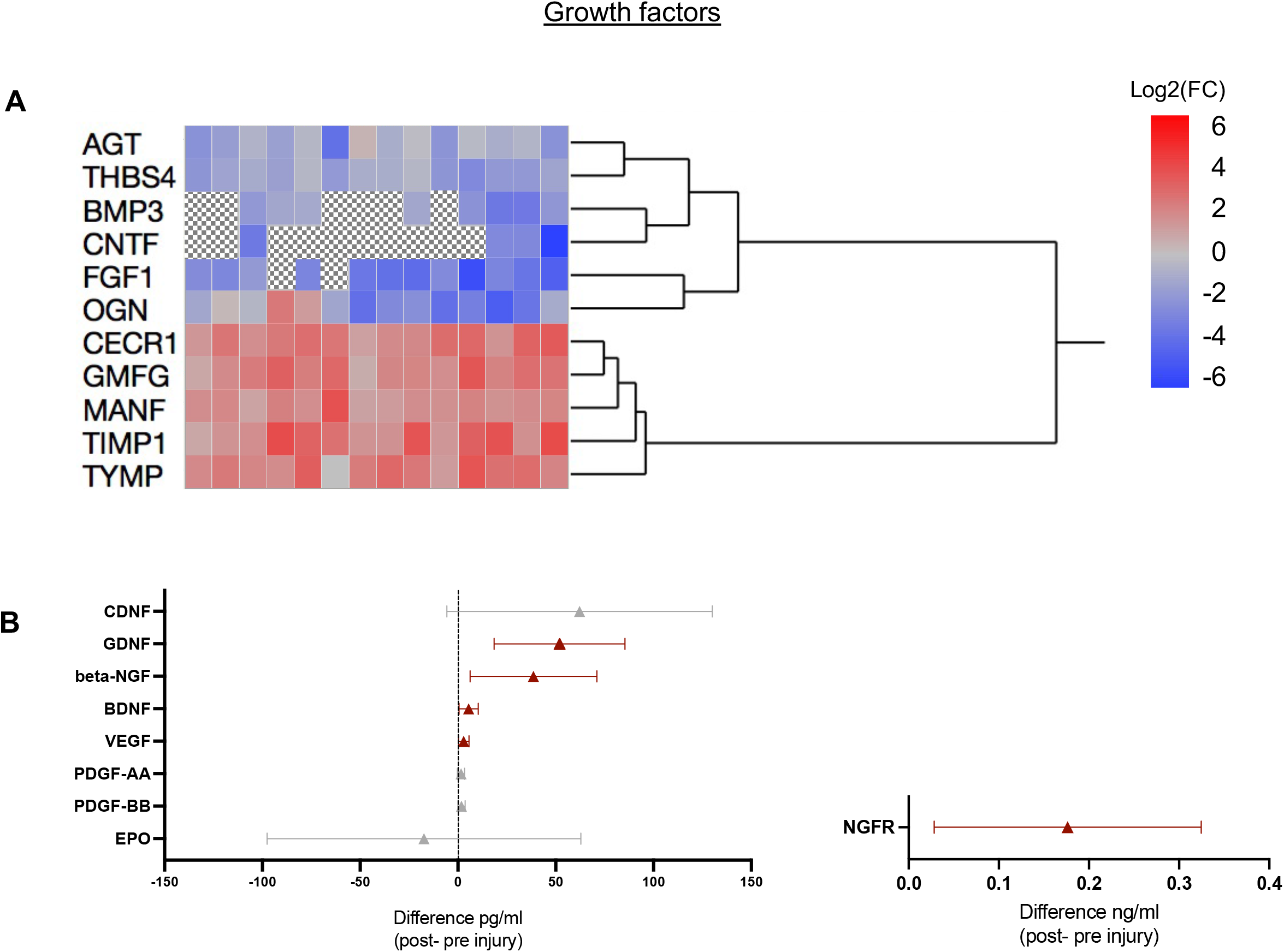
Changes in growth factors after injury based on the GO pathway “Growth Factor Activity. **A.** <u>Mass Spectrometry</u>: Analysis of all the protein products that were significantly upregulated or downregulated (Log2(FC) range: −6 to 6). Proteins are organized by Ward hierarchical clustering. **B.** <u>Immunoassay</u>: Mean (triangle) differences, between naïve and injured tissue samples, and 95% CI for EPO, PDGF-AA and –BB, VEG, BDNF, NGFR, beta-NGF, and GDNF. Red markers represent mean difference CIs that were above zero and gray ones represent mean difference CIs that include zero.

##### Immunoassay

Immunoassays were used to quantify the change in protein content of growth factors of interest that were not abundant enough to be detected via mass spectrometry. These factors of interest included EPO (mean difference: −17.37 pg/ml, n = 13), PDGF-AA (+1.58 pg/ml, n=13) and PDGF–BB (+1.81 pg/ml, n=15), VEGF (+2.96 pg/ml, n=15), BDNF(+5.07 pg/ml, n=15), NGFR (+0.18 ng/ml, n=14), beta-NGF (+38.63 pg/ml n=12), CDNF (62.2 pg/ml n=15), and GDNF (+51.97 pg/ml, n=7), with the 95% confidence interval (CI) of the mean difference between injured and naïve tissues of PDGF-BB (−0.02, 3.65), PDGF-AA (−0.26, 3.42), EPO (−97.6, 62.9), VEGF (0.27, 5.66), BDNF (0.45, 9.68), CDNF (−5.6, 130.1), beta-NGF (6.19, 71.09), NGFR (0.03, 0.32), and GDNF (18.48, 85.45)(Fig 3B).

##### Mass Spectrometry

Fig 4 shows all of the proteins that were significantly upregulated or downregulated from the GO pathway “Growth Factor Binding” (GO: 00019838). Seven proteins were significantly downregulated after transection, and 13 proteins were significantly upregulated after transection. Out of 137 unique genes listed in this GO pathway, 20 gene protein products (15%) were significantly differentially expressed when comparing naïve and injured tissue.

**Fig 4.**
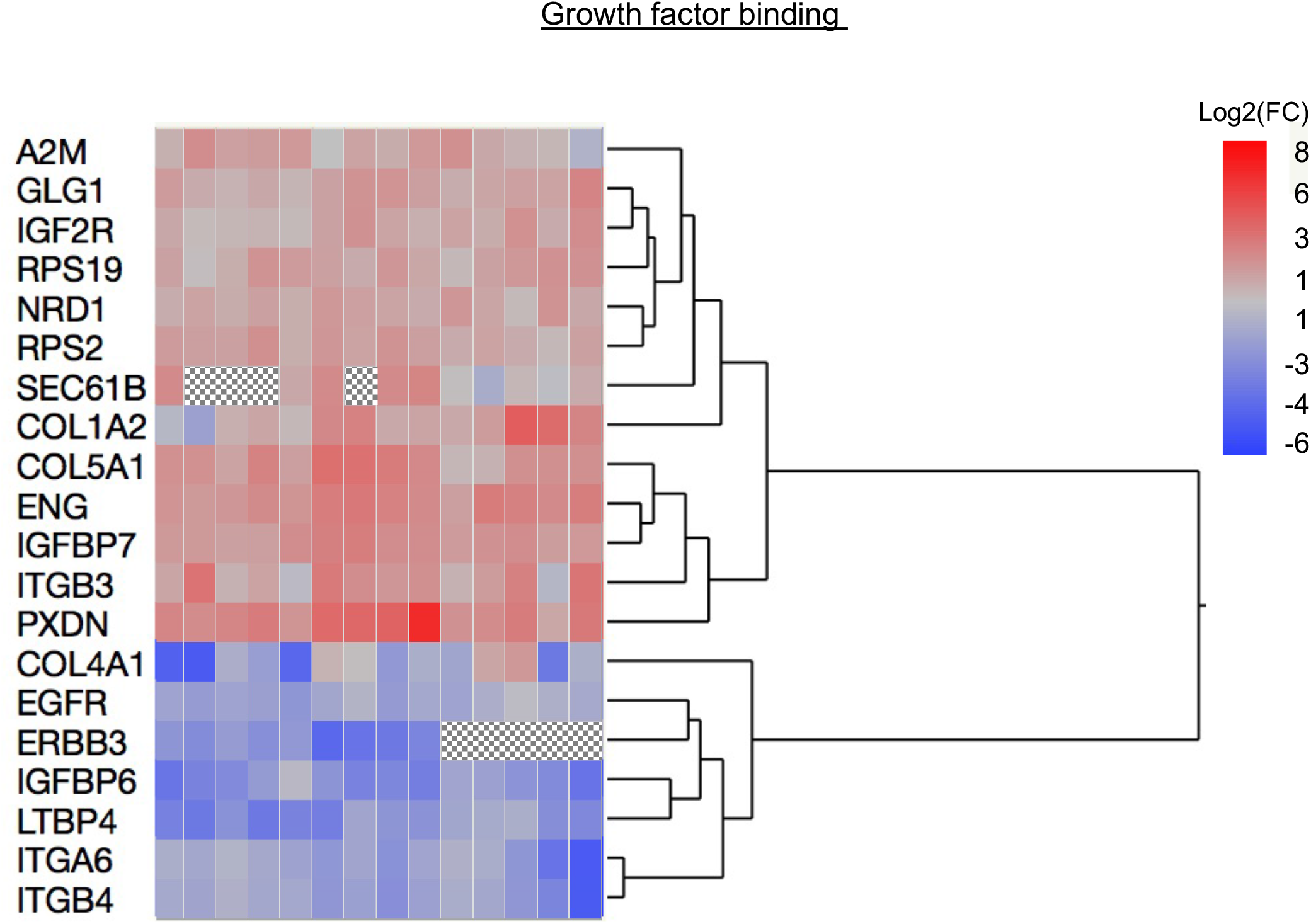
Changes in growth factor binding after injury based on the GO pathway, “Growth Factor Binding.”: Analysis of all the protein products that were significantly upregulated or downregulated from the GO pathway (Log2(FC) range: −6 to 8)

#### Myelination

##### Mass Spectrometry

Fig 5 shows all of the protein products that were significantly upregulated or downregulated from the GO pathway “myelination” (GO: 0042552). Twenty five proteins were significantly downregulated after transection (19% of 132), and 6 proteins (5% of 132) were significantly upregulated after transection. Out of 132 unique genes listed in this GO pathway, 31 gene protein products (24%) were significantly differentially expressed when comparing naïve to injured nerve.

**Fig 5.**
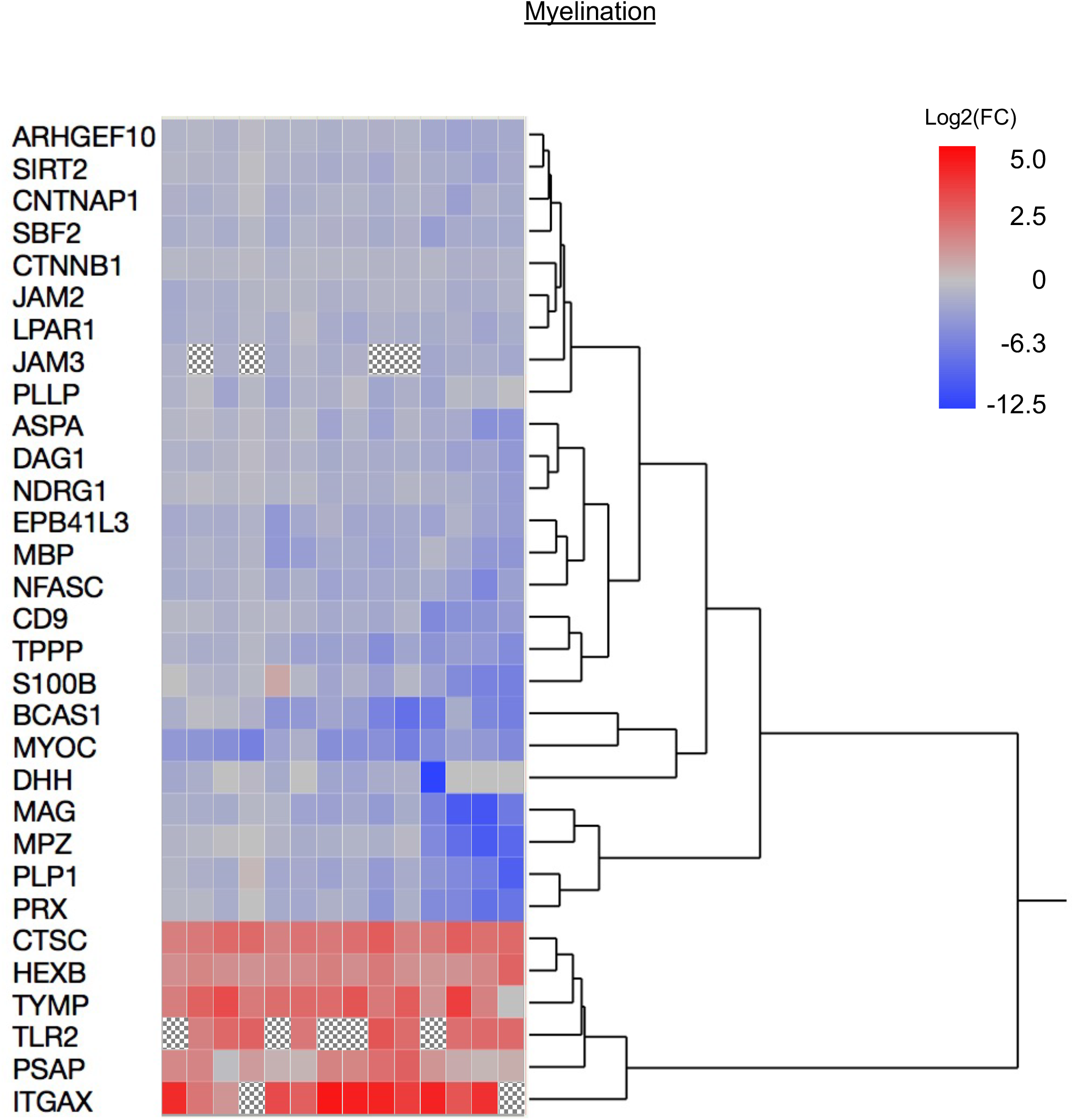
Changes in myelination after injury based on the GO pathway “Myelination.” Analysis of all the protein products that were significantly upregulated or downregulated from the GO pathway (Log2(FC) range: −12.5 to 5)

#### Schwann cell phenotypic changes

##### Schwann Cell Differentiation

###### Mass Spectrometry

Fig 6A shows all of the protein products that were significantly upregulated or downregulated from the GO pathway “Schwann cell differentiation” (GO: 0014037). Ten proteins were significantly downregulated after injury, and no proteins were significantly upregulated after injury. Out of 39 unique genes listed in this GO pathway, 10 gene protein products (26%) were significantly differentially expressed when comparing naïve to injured nerve.

**Fig 6.**
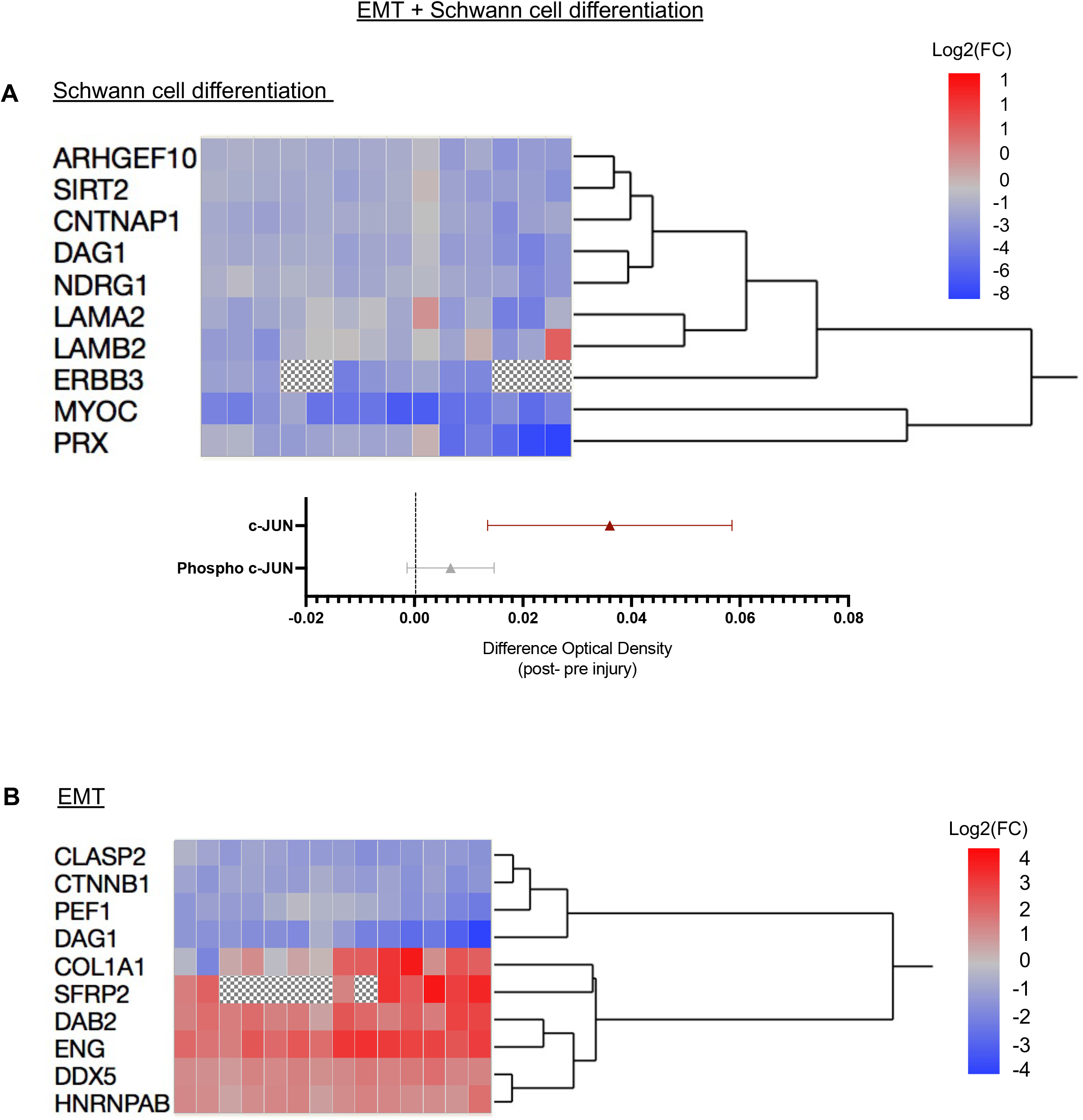
Changes in Schwann cell phenotype after injury based on the GO pathway “Schwann cell differentiation”. **A**. <u>Mass spectrometry</u>: Aall the proteins that were significantly upregulated or downregulated (Log2(FC) range: −4 to 4) **B.** <u>Immunoassay</u>: We quantified the protein expression of c-JUN and its activated form phospho c-JUN, in the naïve and injured tissue and show the mean difference and 95% CI bands. Red markers represent mean difference CIs that were above zero and gray ones represent mean difference CIs that include zero. **C.** Graph of mass spectrometry analysis of all the protein products that were significantly upregulated or downregulated from the GO pathway, “EMT.”

###### Immunoassay

We quantified the content of total c-JUN (mean differerence: +0.03 OD=450, n=15, CI: 0.02, 0.06) and its activated form phospho c-JUN (+0.01 OD=450, n=15, CI: −0.0007, 0.0138) in the naïve and injured tissue(Fig 6A).

##### Epithelial to Mesenchymal Transition (EMT)

###### Mass Spectrometry

Fig 6B shows all of the protein products that were significantly upregulated or downregulated from the GO pathway “EMT” (0001837). Four proteins were significantly downregulated after injury, and 6 proteins were significantly upregulated after injury. Out of 146 unique genes listed in this GO pathway, 10 gene protein products (7%) were significantly differentially expressed when comparing naïve to injured nerve.

#### Anti-apoptosis

##### Mass Spectrometry

Fig 7 shows all of the protein products that were significantly upregulated or downregulated from the pathway “Negative Regulation of Apoptotic Processes” (GO: 0043066). Thirty nine proteins were significantly downregulated after injury, and 42 proteins were significantly upregulated after injury. Out of 890 unique genes listed in this GO pathway, 81 gene protein products (9%) were significantly differentially expressed when comparing naïve to injured nerve. Because we used samples from participants with PD, we particularly examined levels of α-synuclein (SNCA) as aggregates of α-synuclein are associated with PD and deposits of phosporylated α-synuclein[27] are found in the sural nerve of people with PD. We observed a significant reduction (FC: −1.97, *p* = 9.2 x10^-7^) in levels of α-synuclein following injury.

**Fig 7.**
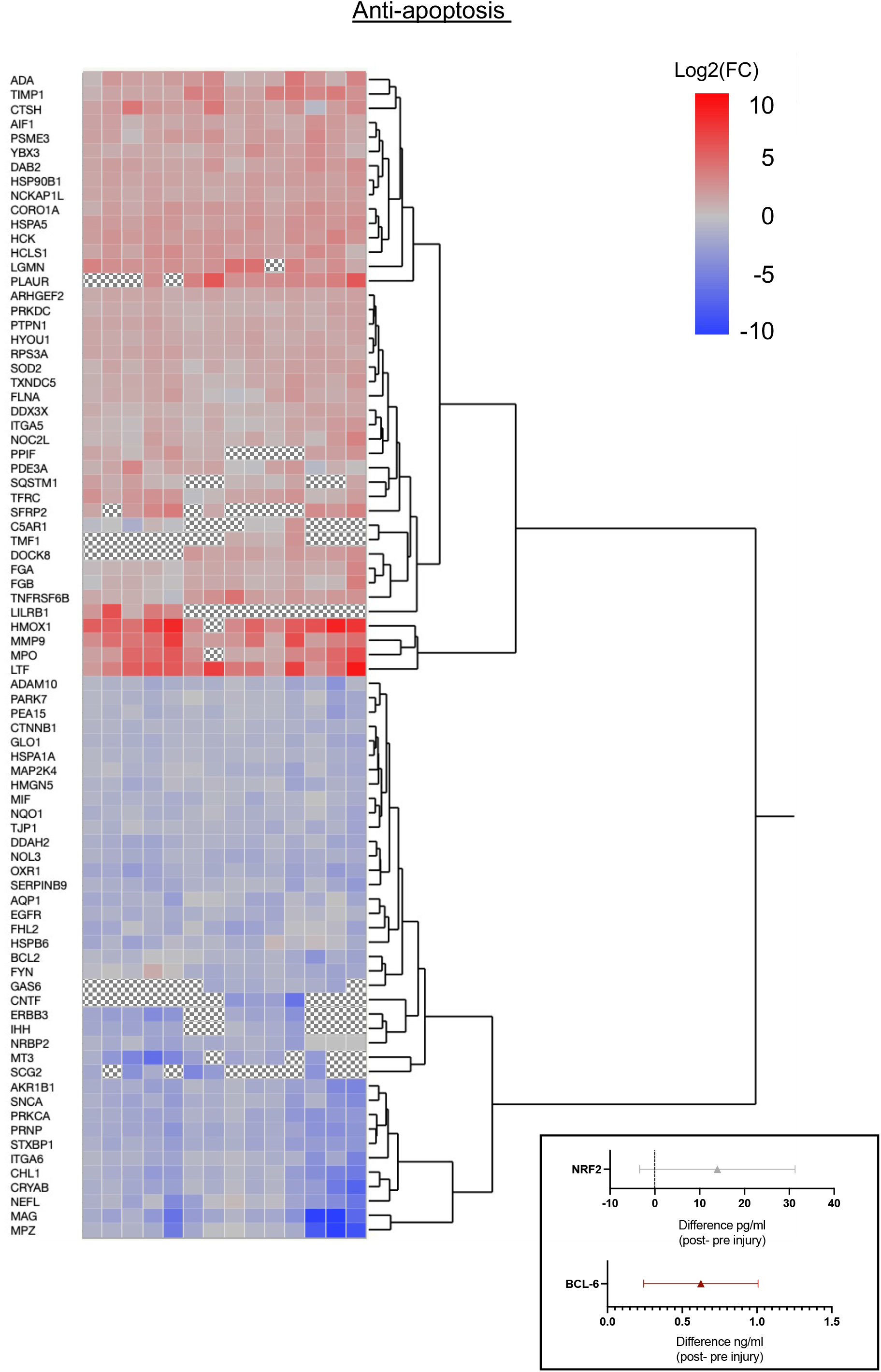
Anti-apoptosis factor changes after injury based on the GO pathway, “Negative Regulation of Apoptotic Processes.” **A**. Mass spectrometry analysis of all the protein products that were significantly upregulated or downregulated (Log2(FC) range: −10 to 10) **B.** <u>Immunoassay</u>: Quantified NRF2 and BCL-6 changes after injury. Red markers represent mean difference CIs that were above zero and gray ones represent mean difference CIs that include zero.

##### Immunoassay

Additionally we examined for NRF2 (mean difference: +13.96 pg/ml, n=14, CI: −3.39, 31.30) and BCL-6 (+0.62 ng/ml, n=15, CI: 0.24, 1.01) where the 95% CI for BCL-6 was above zero in the injured tissue compared to naïve tissue (Fig 7)

#### Response to Axonal Injury

##### Mass Spectrometry

Fig 8A shows all of the protein products that were significantly upregulated or downregulated from the GO pathway, “response to axonal injury” (0048678). This is defined as **“**any process that results in a change in state or activity of a cell or an organism (in terms of movement, secretion, enzyme production, gene expression, etc.) as a result of an axon injury stimulus”. Twelve proteins were significantly downregulated after injury, and 4 proteins were significantly upregulated after injury. Out of 75 unique genes listed in this GO pathway, 16 gene protein products (21%) were significantly differentially expressed when comparing naïve to injured nerve.

**Fig 8.**
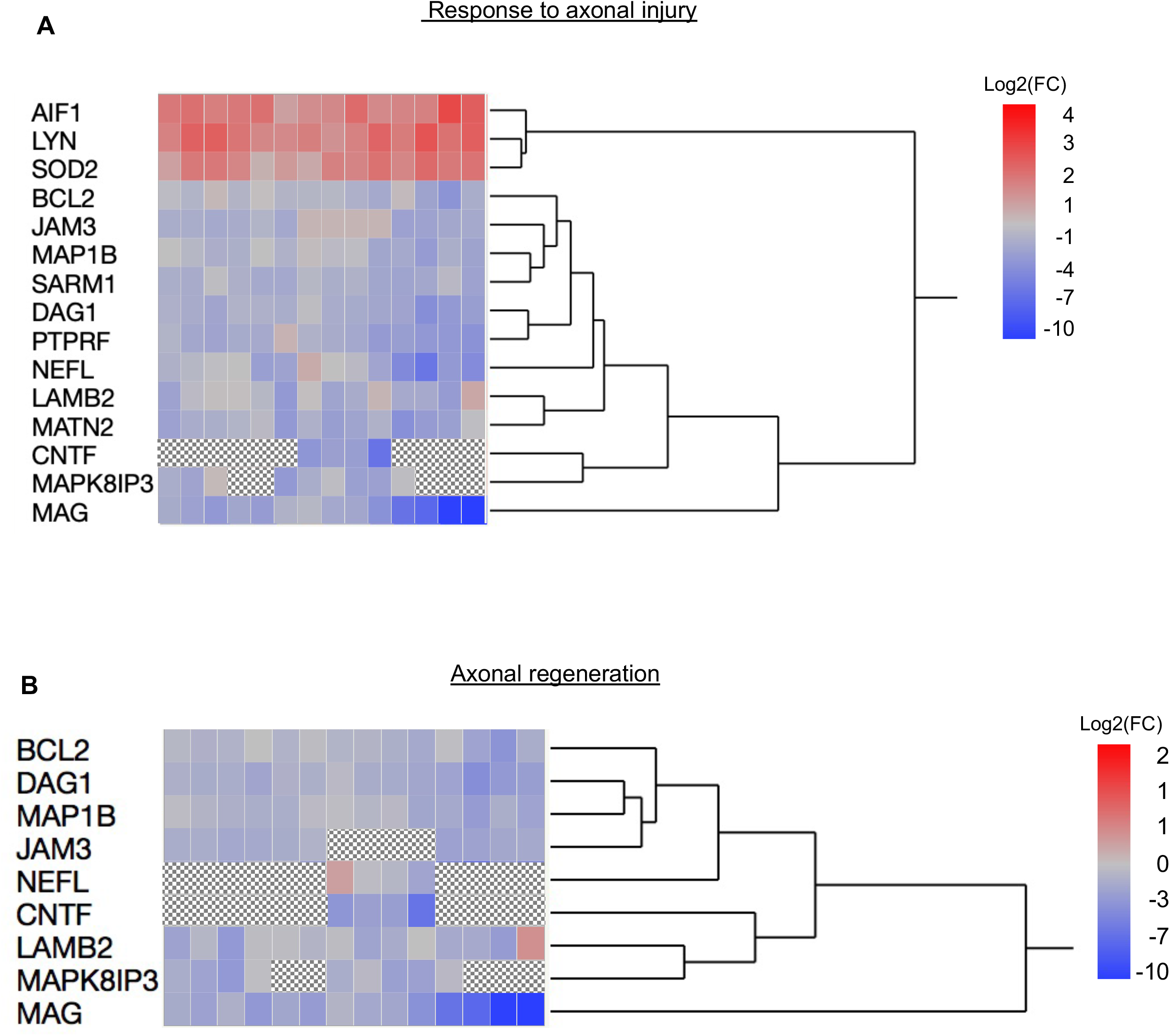
Axonal response after injury. **A.** Graph of mass spectrometry analysis of all the protein products that were significantly upregulated or downregulated from the GO pathway, “Response to Axonal Injury” (Log2(FC) range: −10 to 4) **B.** Graph of mass spectrometry analysis of all the protein products that were significantly upregulated or downregulated from the GO pathway, “Axonal Regeneration” (Log2(FC) range: −10 to 2)

#### Axonal Regeneration

##### Mass Spectrometry

Fig 8B shows all of the protein products that were significantly upregulated or downregulated from the GO pathway, “Axonal regeneration” (0031103). This is defined as **“**The regrowth of axons following their loss or damage.” Nine proteins were significantly downregulated after injury, and one protein was significantly upregulated after injury. Out of 44 unique genes listed in this GO pathway, 10 gene protein products (23%) were significantly differentially expressed when comparing naïve to injured nerve tissue.

## Discussion

The goal of this study was to determine how protein levels change after human peripheral nerve injury. Our study used closely matched internal controls, namely participants’ own naïve and injured peripheral nerve samples to show how the nerve responds with degenerative and regenerative processes. The nerve transection was a necessary event for our underlying trial, an approach which harnesses the regenerative capacity of the peripheral nerve for treating neurodegenerative diseases. Our group was able to transect a peripheral nerve, activate degeneration and regeneration in human sural nerve all *in situ* at the ankle, and collect tissue samples from both conditions, which would not have been ethically possible before this trial. Sural nerve biopsies are common neurosurgical procedures that can result in localized pain and paresthesias[28]. Participants in our studies reported these adverse events acutely but found them tolerable and not bothersome over time. The findings here provide an overview of the proteomic changes in peripheral nerve degeneration/regeneration and could serve as a basis for targeted therapies to promote nerve regeneration and improve clinical outcomes.

### Myelination and schwann cell phenotype after injury

Schwann cells play a major role in supporting peripheral nerves through myelination, maintaining axons, and guiding axons in regeneration after injury. Schwann cells from peripheral nerve tissue have remarkable plasticity in that they rapidly adapt to injury through cellular transformation into a reparative phenotype in two stages. First, upon injury or transection, the transcription factor c-JUN is upregulated, and Schwann cells de-differentiate from a myelinating phenotype into an immature phenotype, undergoing EMT[29,30]. Second, Schwann cells transform into a reparative phenotype and release trophic factors and cytokines to support neuronal survival, and axon regeneration[7,30–32].

Our proteomic results support many of the processes reported in the literature in animal models. Myelin-related proteins such as MPZ, MBP, and MAG are downregulated two weeks after injury (Fig 5)[30]. After injury, the first step of degeneration is clearing the myelin[4]. Schwann cells then de-differentiate. As such, we observed the downregulation of S100B which is expressed by myelinating Schwann cell[33].

c-JUN, is the major transcription factor that activates the Schwann cell repair program by first downregulating the myelinating Schwann cell phenotype[6,7,31,32,34]. Both c-JUN and its activated form, phosphorylated c-JUN, were present more abundantly two weeks after injury than in the naïve tissue (Fig 6A). Interestingly, in our previous work with this transection paradigm[1], *JUN* mRNA levels were not detectable at two weeks suggesting that the peak for mRNA had diminished while the synthesized protein still remained.

c-JUN initiates Schwann cell de-differentiation into a progenitor-like stage, and upregulates stem cell factors such as Sox2, Notch1, Oct6 as a transition between two states. The Schwann cells undergo aEMT-like process in which cells become more like multipotent stem cells, and release neurotrophic factors to support cell survival[2]. *JUN* is rarely expressed in normal, non-injured human nerve tissue[3]; however after injury, it was elevated in human nerve samples denervated for 4–50 days and up to 200 days[29]. In the early response to injury, in many tissues, activation of EMT and stemness is associated with increased cell motility, proliferation, phenotypic flexibility, tissue remodeling and differentiation flexibility[35,36], all which are important in repair and neuroprotection.

Schwann cells transform from the myelinating phenotype to a reparative phenotype via EMT mechanisms[29,30]. Our results show that proteins expressed by Schwann cells such as PRX and LAMA2[37,38] were downregulated after injury. We showed the upregulation of proteins in the EMT GO pathway, including SFRP2 (Fig 6). SFRP2 and SFRP4 are typically associated with concomitant EMT gene expression in a pan-cancer study[39]. The adaptor molecule, disabled-2 (DAB2) promotes EMT[40], and two weeks after injury, the protein was also upregulated in our results[7,30–32] (Fig 6B).

### Growth Factors After Injury

The response to injury creates a neuroprotective microenvironment with the production of neurotrophic factors. Our results here support our hypothesis and previous findings that growth factor levels, specifically neurotrophic factors, are higher after nerve injury[7,30–32]. Two weeks after injury, we observed higher mean levels in the injury samples vs. the naïve samples of growth factors: BDNF, CDNF, VEGF, GDNF, NGF, PDGF-AA and PDGF-BB (Fig 3B). The protein-level profile of individual neuroprotective factors can vary greatly with time. After injury in mice, GDNF peaks around 7 days, and BDNF levels peak around 2–3 weeks[31]. Trophic factors involved in neurite outgrowth, cytoprotection, and neuronal survival such as mesencephalic astrocyte-derived neurotrophic factor (MANF) have also been reported to increase after peripheral nerve injury[41,42]. Interestingly, some growth factors such as CNTF and FGF1 that have been shown in the literature[43–45] to promote nerve regeneration were significantly decreased after the two week injury paradigm, aligning with our findings. This could indicate that the two week timepoint is past the peak expression time for these factors in humans.

Other notable protein level increases from our findings include Glia Maturation Factor Gamma (GMFG) and tissue inhibitor of metalloproteinase-1 (TIMP1). TIMPs play a major role in extracellular matrix (ECM) and tissue remodeling through its action as a potent inhibitor of ECM proteases such as matrix metallopeptidase 9 (MMP9). In a sciatic nerve axotomy model, *Timp1* was found to be the top 6^th^ gene that was induced[46]. Further supporting its role, mice with the *Mmp9* gene knocked out showed greater numbers of de-differentiated/immature myelinating Schwann cells in the injured nerve. This model also suggests that the *Mmp9*/*Timp1*axis guides myelinating Schwann cell differentiation and the molecular assembly of myelin domains during nerve regeneration[46].

### Data Interpretation

Important limitations apply to the interpretation of these results. We used naïve and injured nerve tissue from participants with PD which may differ from non-PD tissue. Synucleinopathies are characteristic in people with PD in both their central and peripheral nervous systems. α-synuclein aggregates have been found in the sciatic nerves and pharyngeal nerves of post-mortem patients with PD[47]. Of note, our results from mass spectrometry show that the α-synuclein protein was significantly downregulated after transection injury. The transection did not cause further upregulation of α-synuclein. Furthermore, as people with PD have a higher incidence of neuropathy[48,49], we recognize that using tissue from participants with PD introduces the concern of neuropathies. The presence of a neuropathy was not an exclusionary criterion for the underlying clinical trial and most participants in the trial did not report a history of neuropathy.

As in Welleford et al.[1], the design of the underlying clinical trial necessitated a delay in flash freezing the samples, more so for the injury samples, that could affect the protein profile. However, the direction of changes in protein levels, after injury, in the GO pathways of myelination, EMT, and axonal regeneration suggest that elements of the proteome in the injured state are in concordance to reported changes in animal studies [2,15,16,31]. The underlying clinical trial design from where these samples were obtained limited our ability to adequately control for the existence of neuropathy, comorbidities that might influence regeneration, age, disease duration, genetics of participants or even the sample freezing time differences among individuals; thus, for interpreting the analyses, we quantified the degree of similarity among different samples by calculating Lin’s CCC (Fig 2B). We found a strong homogeneity among samples within the naïve and injured groups.

Therefore, even with these limitations, we think this study provides insight on the proteomic profile of degenerating/regenerative nerves two weeks after an injury and contributes a database of useful information for establishing biomarkers of peripheral nerve injury in humans.

### Summary

- We present a proteomic picture of the degenerative and regenerative processes in humans after peripheral nerve injury *in situ*
- We provide a database for peripheral nerve repair proteins
- Our results support many processes reported in animal models—down-regulation of myelination, Schwann cell de-differentiation, and upregulation of growth factors

## Acknowledgments

We acknowledge the assistance of IDeA National Resource for Quantitative Proteomics at UAMS and Drs. Sam Mackintosh and Stephanie Byrum. The IDeA National Resource for Quantitative Proteomics at UAMS is supported by NIH grant R24GM137786. We would also like to acknowledge the Research Flow Cytometry Core in the Division of Rheumatology at Cincinnati Children’s Hospital Medical Center and Alyssa Sproles. We thank Morgan Yazell for trial execution, Tom Dolan for medical illustration, Drs. Randal Voss and Jeramiah Smith for guidance on GO analyses and heatmaps, and Dr. Andrew Welleford for analysis guidance.

## Institutional Review Board Statement

The University of Kentucky’s Institutional Review Board approved the study (44749).

## Informed Consent

The participants provided written informed consent.

## References

1. Welleford AS, Quintero JE, Seblani NE, Blalock E, Gunewardena S, Shapiro SM, et al. RNA Sequencing of Human Peripheral Nerve in Response to Injury: Distinctive Analysis of the Nerve Repair Pathways. Cell Transplant. 2020;29:963689720926157.

2. Clements MP, Byrne E, Camarillo Guerrero LF, Cattin AL, Zakka L, Ashraf A, et al. The Wound Microenvironment Reprograms Schwann Cells to Invasive Mesenchymal-like Cells to Drive Peripheral Nerve Regeneration. Neuron. 2017;96(1):98–114 e7.

3. Hutton EJ, Carty L, Laura M, Houlden H, Lunn MP, Brandner S, et al. c-Jun expression in human neuropathies: a pilot study. J Peripher Nerv Syst. 2011;16(4):295–303.

4. Pellegrino RG, Politis MJ, Ritchie JM, Spencer PS. Events in degenerating cat peripheral nerve: induction of Schwann cell S phase and its relation to nerve fibre degeneration. J Neurocytol. 1986;15(1):17–28.

5. Arthur-Farraj PJ, Latouche M, Wilton DK, Quintes S, Chabrol E, Banerjee A, et al. c-Jun reprograms Schwann cells of injured nerves to generate a repair cell essential for regeneration. Neuron. 2012;75(4):633–47.

6. Fontana X, Hristova M, Da Costa C, Patodia S, Thei L, Makwana M, et al. c-Jun in Schwann cells promotes axonal regeneration and motoneuron survival via paracrine signaling. J Cell Biol. 2012;198(1):127–41.

7. Jessen KR, Mirsky R. The repair Schwann cell and its function in regenerating nerves. J Physiol. 2016;594(13):3521–31.

8. La Fleur M, Underwood JL, Rappolee DA, Werb Z. Basement membrane and repair of injury to peripheral nerve: defining a potential role for macrophages, matrix metalloproteinases, and tissue inhibitor of metalloproteinases-1. J Exp Med. 1996;184(6):2311–26.

9. Mueller M, Leonhard C, Wacker K, Ringelstein EB, Okabe M, Hickey WF, et al. Macrophage response to peripheral nerve injury: the quantitative contribution of resident and hematogenous macrophages. Lab Invest. 2003;83(2):175–85.

10. Stratton JA, Holmes A, Rosin NL, Sinha S, Vohra M, Burma NE, et al. Macrophages Regulate Schwann Cell Maturation after Nerve Injury. Cell Rep. 2018;24(10):2561–72 e6.

11. Sebille A, Bondoux-Jahan M. Motor function recovery after axotomy: enhancement by cyclophosphamide and spermine in rat. Exp Neurol. 1980;70(3):507–15.

12. Allodi I, Udina E, Navarro X. Specificity of peripheral nerve regeneration: interactions at the axon level. Prog Neurobiol. 2012;98(1):16–37.

13. Blesch A, Lu P, Tsukada S, Alto LT, Roet K, Coppola G, et al. Conditioning lesions before or after spinal cord injury recruit broad genetic mechanisms that sustain axonal regeneration: superiority to camp-mediated effects. Exp Neurol. 2012;235(1):162–73.

14. Nutt JG, Burchiel KJ, Comella CL, Jankovic J, Lang AE, Laws ER, Jr., et al. Randomized, double-blind trial of glial cell line-derived neurotrophic factor (GDNF) in PD. Neurology. 2003;60(1):69–73.

15. Naveilhan P, ElShamy WM, Ernfors P. Differential regulation of mRNAs for GDNF and its receptors Ret and GDNFR alpha after sciatic nerve lesion in the mouse. Eur J Neurosci. 1997;9(7):1450–60.

16. Heumann R, Korsching S, Bandtlow C, Thoenen H. Changes of nerve growth factor synthesis in nonneuronal cells in response to sciatic nerve transection. J Cell Biol. 1987;104(6):1623–31.

17. Brushart TM, Aspalter M, Griffin JW, Redett R, Hameed H, Zhou C, et al. Schwann cell phenotype is regulated by axon modality and central-peripheral location, and persists in vitro. Exp Neurol. 2013;247:272–81.

18. Meyer M, Matsuoka I, Wetmore C, Olson L, Thoenen H. Enhanced synthesis of brain-derived neurotrophic factor in the lesioned peripheral nerve: different mechanisms are responsible for the regulation of BDNF and NGF mRNA. J Cell Biol. 1992;119(1):45–54.

19. Gonzalez A, Ugarte G, Restrepo C, Herrera G, Pina R, Gomez-Sanchez JA, et al. Role of the Excitability Brake Potassium Current IKD in Cold Allodynia Induced by Chronic Peripheral Nerve Injury. J Neurosci. 2017;37(12):3109–26.

20. Boyd JG, Gordon T. Glial cell line-derived neurotrophic factor and brain-derived neurotrophic factor sustain the axonal regeneration of chronically axotomized motoneurons in vivo. Exp Neurol. 2003;183(2):610–9.

21. Scheib J, Hoke A. Advances in peripheral nerve regeneration. Nat Rev Neurol. 2013;9(12):668–76.

22. Grothe C, Haastert K, Jungnickel J. Physiological function and putative therapeutic impact of the FGF-2 system in peripheral nerve regeneration--lessons from in vivo studies in mice and rats. Brain Res Rev. 2006;51(2):293–9.

23. van Horne CG, Quintero JE, Gurwell JA, Wagner RP, Slevin JT, Gerhardt GA. Implantation of autologous peripheral nerve grafts into the substantia nigra of subjects with idiopathic Parkinson’s disease treated with bilateral STN DBS: a report of safety and feasibility. J Neurosurg. 2017;126(4):1140–7.

24. van Horne CG, Quintero JE, Slevin JT, Anderson-Mooney A, Gurwell JA, Welleford AS, et al. Peripheral nerve grafts implanted into the substantia nigra in patients with Parkinson’s disease during deep brain stimulation surgery: 1-year follow-up study of safety, feasibility, and clinical outcome. J Neurosurg. 2018;129(6):1550–61.

25. Taylor SL, Leiserowitz GS, Kim K. Accounting for undetected compounds in statistical analyses of mass spectrometry ‘omic studies. Stat Appl Genet Mol Biol. 2013;12(6):703–22.

26. Wei R, Wang J, Su M, Jia E, Chen S, Chen T, et al. Missing Value Imputation Approach for Mass Spectrometry-based Metabolomics Data. Sci Rep. 2018;8(1):663.

27. Zhang J, Li X, Li JD. The Roles of Post-translational Modifications on alpha-Synuclein in the Pathogenesis of Parkinson’s Diseases. Front Neurosci. 2019;13:381.

28. Hilton DA, Jacob J, Househam L, Tengah C. Complications following sural and peroneal nerve biopsies. J Neurol Neurosurg Psychiatry. 2007;78(11):1271–2.

29. Wilcox MB, Laranjeira SG, Eriksson TM, Jessen KR, Mirsky R, Quick TJ, et al. Characterising cellular and molecular features of human peripheral nerve degeneration. Acta Neuropathol Commun. 2020;8(1):51.

30. Jessen KR, Mirsky R. The Success and Failure of the Schwann Cell Response to Nerve Injury. Front Cell Neurosci. 2019;13:33.

31. Jessen KR, Arthur-Farraj P. Repair Schwann cell update: Adaptive reprogramming, EMT, and stemness in regenerating nerves. Glia. 2019;67(3):421–37.

32. Jessen KR, Mirsky R, Lloyd AC. Schwann Cells: Development and Role in Nerve Repair. Cold Spring Harb Perspect Biol. 2015;7(7):a020487.

33. Mata M, Alessi D, Fink DJ. S100 is preferentially distributed in myelin-forming Schwann cells. J Neurocytol. 1990;19(3):432–42.

34. Parkinson DB, Bhaskaran A, Arthur-Farraj P, Noon LA, Woodhoo A, Lloyd AC, et al. c-Jun is a negative regulator of myelination. J Cell Biol. 2008;181(4):625–37.

35. Nieto MA, Huang RY, Jackson RA, Thiery JP. Emt: 2016. Cell. 2016;166(1):21–45.

36. Forte E, Chimenti I, Rosa P, Angelini F, Pagano F, Calogero A, et al. EMT/MET at the Crossroad of Stemness, Regeneration and Oncogenesis: The Ying-Yang Equilibrium Recapitulated in Cell Spheroids. Cancers (Basel). 2017;9(8).

37. Gillespie CS, Lee M, Fantes JF, Brophy PJ. The gene encoding the Schwann cell protein periaxin localizes on mouse chromosome 7 (Prx). Genomics. 1997;41(2):297–8.

38. Previtali SC, Zambon AA. LAMA2 Neuropathies: Human Findings and Pathomechanisms From Mouse Models. Front Mol Neurosci. 2020;13:60.

39. Vincent KM, Postovit LM. A pan-cancer analysis of secreted Frizzled-related proteins: re-examining their proposed tumour suppressive function. Sci Rep. 2017;7:42719.

40. Prunier C, Howe PH. Disabled-2 (Dab2) is required for transforming growth factor beta-induced epithelial to mesenchymal transition (EMT). J Biol Chem. 2005;280(17):17540–8.

41. Eesmaa A, Yu LY, Goos H, Noges K, Kovaleva V, Hellman M, et al. The cytoprotective protein MANF promotes neuronal survival independently from its role as a GRP78 cofactor. J Biol Chem. 2021:100295.

42. Wen W, Wang Y, Li H, Xu H, Xu M, Frank JA, et al. Mesencephalic Astrocyte-Derived Neurotrophic Factor (MANF) Regulates Neurite Outgrowth Through the Activation of Akt/mTOR and Erk/mTOR Signaling Pathways. Front Mol Neurosci. 2020;13:560020.

43. Sahenk Z, Seharaseyon J, Mendell JR. CNTF potentiates peripheral nerve regeneration. Brain Res. 1994;655(1-2):246–50.

44. Hu Y, Leaver SG, Plant GW, Hendriks WT, Niclou SP, Verhaagen J, et al. Lentiviral-mediated transfer of CNTF to schwann cells within reconstructed peripheral nerve grafts enhances adult retinal ganglion cell survival and axonal regeneration. Mol Ther. 2005;11(6):906–15.

45. Midha R, Munro CA, Dalton PD, Tator CH, Shoichet MS. Growth factor enhancement of peripheral nerve regeneration through a novel synthetic hydrogel tube. J Neurosurg. 2003;99(3):555–65.

46. Kim Y, Remacle AG, Chernov AV, Liu H, Shubayev I, Lai C, et al. The MMP-9/TIMP-1 axis controls the status of differentiation and function of myelin-forming Schwann cells in nerve regeneration. PLoS One. 2012;7(3):e33664.

47. Mu L, Sobotka S, Chen J, Su H, Sanders I, Adler CH, et al. Alpha-synuclein pathology and axonal degeneration of the peripheral motor nerves innervating pharyngeal muscles in Parkinson disease. J Neuropathol Exp Neurol. 2013;72(2):119–29.

48. Grambalova Z, Kaiserova M, Vastik M, Mensikova K, Otruba P, Zapletalova J, et al. Peripheral neuropathy in Parkinson’s disease. Neuro Endocrinol Lett. 2015;36(4):363–7.

49. Zis P, Grunewald RA, Chaudhuri RK, Hadjivassiliou M. Peripheral neuropathy in idiopathic Parkinson’s disease: A systematic review. J Neurol Sci. 2017;378:204–9.

